# Impact of Propionic Acidemia on Brain Astrocytes

**DOI:** 10.1101/2021.02.07.428966

**Authors:** Maria L Cotrina, Wei Sun, Michael Chen, Adam J Guenzel, José Zhagnay, Michael A Barry, James Goldman, Maiken Nedergaard

## Abstract

Propionic acidemia (PA) is an inborn error of metabolism (IEM) caused by mutations in the enzyme propionyl CoA carboxylase (PCC). It is characterized by the inability to break down branched chain amino acids and odd chain fatty acids, causing a buildup of toxic organic acids in blood. PA affects every organ in the body with particularly severe manifestations in the brain, like hyperammonemia, hypomyelination, seizures, cognitive impairments, optic nerve atrophy and autism spectrum disorders. Dietary management and liver transplantation have helped to ameliorate the acute expression of the disorder, but do not prevent the chronic toxicity that builds up in brain. Despite the severe brain manifestation of the disease, little is known about the mechanisms by which PA affects the nervous system.

PCCA and PCCB, the two subunits required for a functional PCC enzyme, are both expressed not only in neurons but also in astrocytes. Using the two rodent genetic models of PA currently available, with mutations in PCCA, we have evaluated the involvement of astrocytes in the neuropathology of propionic acidemia. These mice exhibit cardiac pathology and hyperammonemia, similar to what is observed in patients with PA.

We found that wild type (wt) astrocytes positively respond to treatment with L-Carnitine, a therapeutic approach commonly used in patients with PA, by improved survival and more efficient mitochondrial morphology. Transcriptome analysis from astrocytes derived from the wt or the mutant mice confirm that these astrocytes lack exons 3 and 4 like in the human mutations of PA. However, no other genes/exons were statistically significant with regards to differential expression between astrocytes derived from KO or from WT animals, suggesting that astrocytes in culture may be able to compensate the PCC deficiency.

Histological analysis of neuronal and glial markers during brain development (TUJ1, MAP2 for neurons; nestin and Iba1 for glia) do not show significant alterations neither in distribution nor numbers of cells in the developing brain of the PCCA-/- mice. Analysis in the adult brain of mutant mice shows some variable degree of microgliosis but no indication of reactive astrocytes. No gross abnormalities were observed in cortex, hippocampus, striatum or cerebellum of adult brains of PCCA-/- mice, either.

In summary, astrocytes from PCCA deficient mice show surprisingly little alterations both *in vitro* and *in vivo*. Our results evidence the need to further understand the effects of PA in brain cells to help develop potential new therapies that can preserve brain function in children affected by this devastating disease.

**HIGHLIGHTS:**

- Astrocytes detoxify ammonia in brain and may be affected by propionic acidemia, a disorder that causes hyperammonemia in brain.
- RNA seq of astrocytes in culture derived from PCCA mutant mice does not show effect in these cells.
- *In vivo* analysis of glial and neuronal cells also shows no difference in development or adult mutant mice
- Astrocytes may not be an adequate target of clinical therapies in this disorder

## INTRODUCTION

Metabolic diseases affect every organ in the body, but they have particularly noxious effects in the brain. In patients with propionic acidemia (PA), for example, neural manifestations of the disorder include hypotonia, hyperammonemia, hypomyelination, seizures, cognitive impairments and autism spectrum disorders (Van Gosen, 2008; Schreiber et al., 2012, Cotrina et al., 2019). A neurological, non-metabolic manifestation of the disease has also been reported (Nyhan et al., 1999; Almuqbil et al., 2019).

A permanent risk of acute metabolic stroke exists even with a relatively good control of blood propionic acid levels (Haas et al., 1995; Kalidas and Behrouz, 2008; Scholl-Bürgi et al., 2009), suggesting that brain metabolism manage the build-up of these toxic acids differently from and independently of other organs.

The mechanism by which PA causes brain pathology is unknown. The “toxic-metabolite hypothesis” sustains that propionyl-CoA and 2-methycitrate are directly responsible for neurotoxicity (Ballhausen et al., 2009). Neurodegeneration could also be caused by the production of high levels of dicarboxylic acids that overload the capacity of the corresponding dicarboxylate transporters (Kölker et al., 2006; Sauer et al., 2006) resulting in slow clearance through the blood-brain barrier (the so-called “trapping hypothesis”). Last, ammonia is chronically elevated in children with PA (Surtees et al., 1992; Filipowicz et al., 2006), but it is not clear what role it plays in brain pathology in this disease.

There are currently no effective treatments to prevent the neurological deterioration observed in PA patients.

Neurodegenerative diseases have long been thought to result exclusively from the loss of neuronal cells. But many, if not all, of these conditions have been recently found to show a surprisingly high degree of astrocytic involvement like in diseases of aging (Alzheimer’s, Parkinson’s and Huntington’s diseases) and a wide array of neurological syndromes, (ischemia, depression and epilepsy). Even demyelinating leukodystrophies like Alexander’s disease (the only known human disease solely caused by mutations in the astrocytic protein GFAP), Vanishing White Matter and cerebellar ataxia, show impaired astrocyte function of astrocytes (Allaman et al., 2011; Dietrich et al., 2005; Bugiani et al., 2010).

Whether the mutated enzyme in PA, propionyl CoA carboxylase (PCC) is located in neurons or astrocytes or both is still under debate. For example, the presence of both mRNA and protein for propionyl CoA carboxylase alpha (PCCα) has been documented in embryonic and adult rat CNS, with highest levels in neurons and no expression in other brain cells, like oligodendrocytes or astrocytes (Ballhausen et al., 2009). This result contrasts with a previous study that found PCC enzymatic activity in primary neuronal and glial cell cultures and cell lines of glial origin (Suormala et al., 2002). Nguyen et al. (2007) have also reported PCC activity in glial cells in vivo and in vitro; 3-C14-propionate increases specific activity of glutamine by several fold compared to glutamate, suggesting that propionate is metabolized via the TCA-cycle in astrocytes. Propionic acid has also been shown to impact astrocytic cytoskeletal proteins in vitro (Vieira et al., 2006).

The presence of PCC in astrocytes, therefore, is still highly debatable, with antibody data supporting its sole presence in neurons but enzymatic data suggesting its activity in astrocytes.

Regardless of the direct expression and function of PCC in astrocytes, these glial cells may be very vulnerable to PA because of their direct contact with blood vessels. Ammonia can diffuse through cell membranes (Bosoi and Rose, 2009) and in astrocytes is converted into glutamine. Indeed, astrocytes, but not neurons, contain glutamine synthetase and are thus the main neural cell type with the capacity to detoxify the high ammonia levels that accumulate in brain of PA patients (Albrecht et al., 2010). Also, abnormal accumulation of secondary metabolites from branched chain amino acids (BCAAs)—the amino acids that PA patients cannot metabolize correctly—is highest at the level of the blood-brain barrier (Knerr et al., 2010), precisely where astrocytic end-feet are located.

Thus, independently of the direct presence of PCC in glia, astrocytes are likely to be affected by PA because they are the first to receive excess ammonia and secondary toxic metabolites from blood, owing to their highly abundant contacts with blood vessels.

In this study, we investigate the contribution of glial cells to the PA phenotype in brain utilizing the PA mouse models currently available (Miyazaki et al., 2001; Guenzel et al., 2013).

To our knowledge, this is the first study to address the involvement of astrocytes in PA in an animal model. This information is critical to design better therapies that protect this organ in children with PA.

## METHODS

### Animals

PCCA-/- (Miyazaki et al., 2001) or wild type mice were bred from heterozygous PA+/- parents and sacrificed at P0-P2. Genotype was confirmed by PCR as described in Hofherr et al (2009). Mice were housed in the animal facility of the University of Rochester with a 12-h light/dark cycle and allowed water and food *ad libitum*. For experiments with the hypomorphic model of PA (Guenzel et al., 2013), mice were raised in the animal facility of the Mayo Clinic and genotyped as described. Two wt/het adult animals and 2 PCCA-/- (A138T mutant) of the same litter were used from these hypomorphic mice.

#### RNA sequencing of astrocytes mRNA

Cultured astrocytes derived from 4 PCCA-/- and 4 wild-type (WT) P0 animals were subject to mRNA isolation and sequencing using Illumina RNA-Seq at the UR Genomics Research Center. Realignments were done using STAR and DEXSeq/R was used for DE analysis and visualization. The threshold set up for statistical significance throughout was p<0.05. To confirm the results obtained from the first analysis, two subsequent analyses were done with TopHat 2.0 and SHRiMP, for alignment readout.

#### Astrocyte culture and exposure to H_2_O_2_ and carnitine

Pure astrocytic cultures were prepared from P0-P2 animals as in Cotrina et al. (1998a) and Nedergaard (1995). 10-15 days old astrocytic cultures were subject to a sublethal dose of H_2_O_2_ (0.7-1.5 mM) for 45 minutes and subsequently incubated in absence or presence of 500 uM L-carnitine (Sigma-Tau) for 2 hours. After washes in PBS, cultures were stained with the cell-death indicator propidium iodide (PI).

For mitochondrial morphology studies, cultures were subsequently incubated with mito-tracker Red CMXRos probe for 10 min at 37C (Molecular Probes). Cultures were then fixed in 4% paraformaldehyde (PFA), rinsed in PBS and kept at 4C° until imaging.

For immunocytochemistry, cultures were fixed with 4% PFA after carnitine incubation, rinsed in PBS, incubated with anti-GFAP antibodies and counterstained with 647 or 488 Alexa-antibodies (Molecular Probes).

### Immunohistochemistry

Standard immunohistochemistry protocols were performed as in Cotrina et al. (2008a). For staining in P0-P2 animals, the brains were fixed by immersion o/n in 4% paraformaldehyde (PFA) followed by two rinses in PBS. Brains were kept at 4C° until use, no later than a week after fixation. For studies in the adult mice, the animals were perfused with 4% PFA, dissected out and left o/n in PFA. The protocol after dissection was the same than for the neonatal brains.

The antibodies used were: GFAP for astrocytes (Chemicon or Dako 1:500), GS for astroyctes (Chemicon, 1:500), MAP2 (AbCAM 1:5000) and Tuj1 (Covance, 1:500) for neurons, Iba1 for microglia (Waco, 1:500), Cx43 for astrocytes (Chemicon 1:500), AQP4 for astrocytes (Alomone 1:500).

## RESULTS

### Transcriptome of PA astrocytes

To date, no information exists about how ions and metabolites are exchanged between neurons and astrocytes under PA stress conditions.

Our first set of experiments aimed at screening for genes that are expressed differently in astrocytes of PCCA-/- knockout (KO) mouse compared to wt mice. To this end, we set up, mRNA sequencing to analyze all transcripts in the samples.

Cultured astrocytes derived from 4 KO and 4 wild-type (WT) P0 animals were subject to mRNA isolation and sequencing using Illumina RNA-Seq at the UR Genomics Research Center. Realignments were done using STAR and DEXSeq/R was used for DE analysis and visualization. The threshold set up for statistical significance throughout was p<0.05. To confirm the results obtained from the first analysis, two subsequent analyses were done with TopHat 2.0 and SHRiMP, for alignment readout.

From the genome viewer, we confirmed that PCCA exons 3 and 4 were not expressed in any of the KO samples. The rest of the exons were expressed in the KO and all the exons were expressed in the WT. From all the analyses utilized, we found that no other genes/exons were statistically significant with regards to differential expression. This means that we found no difference between astrocytes derived from KO or from WT animals (**Figure 1**).

**Figure 1.**
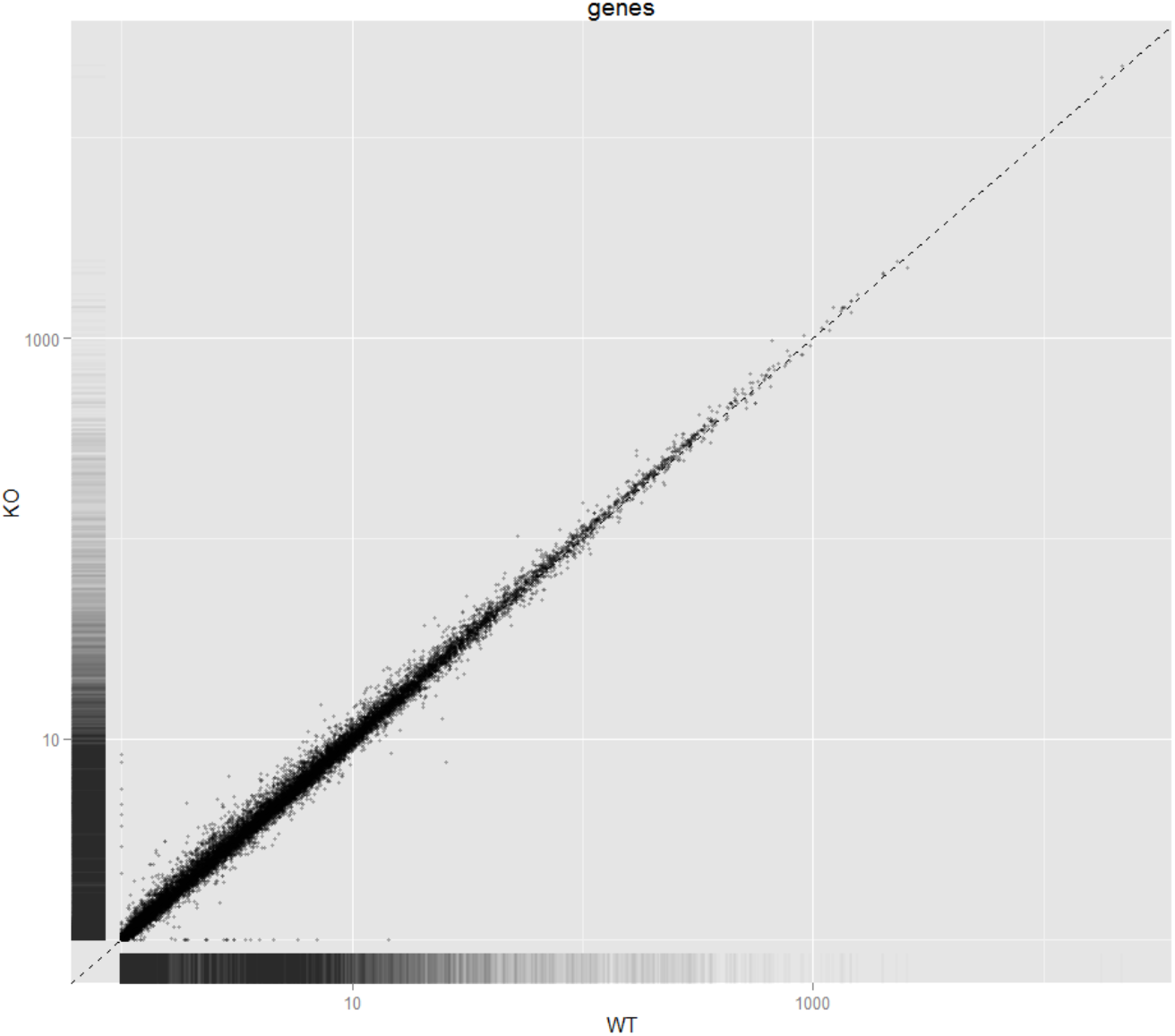
Expression Scatterplot for genes in the knock-out vs wild type astrocytes. The distance of each dot from the diagonal line is directly proportional to the differential expression between the two phenotypes.

### PA does not affect brain development in the neonatal P0 PCCA-/- mice

Given the results obtained from the mRNA seq analysis, we next focused on the characterization of the brain pathology caused by PA in the neonatal PCAA-/- mouse *in vivo*.

We analyzed two P0 neonatal litters containing, at least, 2 KO and 2-3 heterozygote/wild-type mice in each. Whereas some of the other brain pathologies we have studied in our laboratory promote substantial delays in brain development, we could not find any indication that the brain of P0 PCCA-/- mice develops differently from that of the WT brain. **Figure 2** shows representative results of this part of the study where two different markers of neuronal development are used at the same time: on the one hand, TUJ1, a neuronal axonal marker that is expressed from early neuronal development and, on the other hand, MAP2, another neuronal marker that appears slightly later in development, as neurons mature fully. Normally, the merger of the two colors (green for TUJ and red for MAP2) displays as orange. However, if there were a delay in development in the KO mice, we would expect a higher proportion of the green TUJ1 signal (the early marker) with respect to the later red marker MAP2. As observed in Figure 2, no significant differences were observed in any of the areas analyzed in the P0 brains.

**Figure 2.**
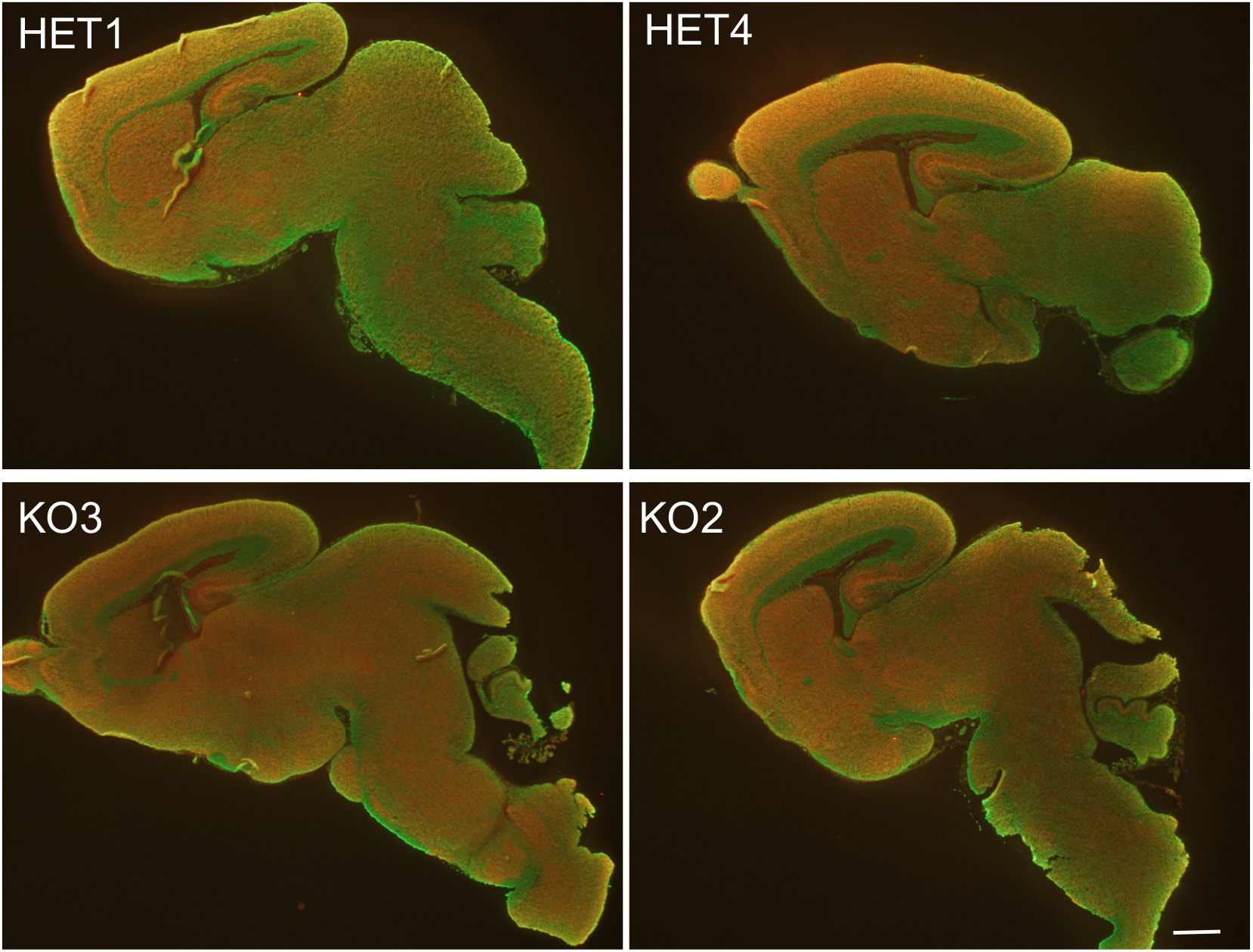
Neural development is unaltered in PCCA -/- mice. Representative staining of TUJ1/MAP2 (green/red) double staining in P0 mice brains to detect differences in the rate of neural development shows that no overall changes occur in k/o mice (KO3 and KO2) compared to het mice (HET1 and HET4) as the merge of the two colors is similar. No gross abnormalities are observed in any of the areas analyzed. Scale bar, 500 μm

Because at P0 there are no GFAP+ cells that correspond to mature astrocytes and almost no myelinating cells (oligodendrocytes), we then stained our PCCA-/- sections with antibodies against nestin, a marker for progenitor cells (NPCs) that is later replaced by specific neuronal and glial proteins. Consistent with the data obtained above, we also observed normal glial development, as nestin+ cells were equally distributed in both KO and heterozygous/WT animals (**Figure 3**).

**Figure 3.**
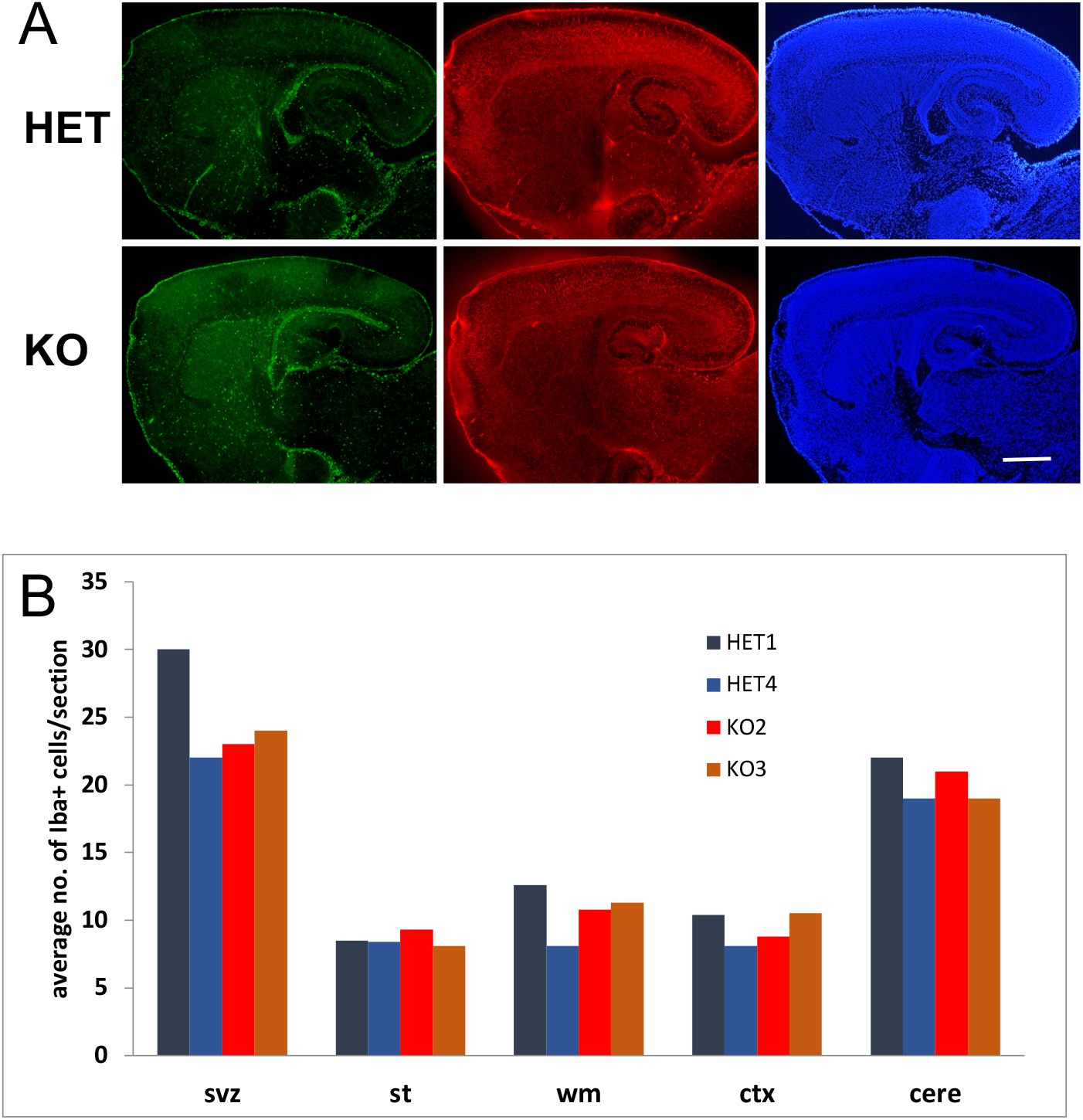
Glial development is normal in PCCA -/- mice. Iba1 (green) and nestin (red) levels in k/o and het P0 mice. No differences were observed either in the distribution (A, green) or the number of cells (graph in B) of the microglial lineage (Iba+ cells) or the astrocyte/oligodendroglial lineage (A, red; nestin+ cells). Sections are counterstained with DAPI for nuclear labeling (blue) Scale bar, 500 μm. **svz**, subventricular zone; **st**, striatum; **wm**, white matter; **ctx,** cortex; **cere**, cerebellum

We also wanted to investigate whether proliferation and migration of the other glial cell type of the brain, the microglia, also proceeded normally. Although microglial cells are usually associated with responses to injury in the brain, they are critical for proper synaptic formation during development (reviewed in Liddelow et al., 2020). We thus stained P0 brain sections with the microglial marker Iba1, specific for this cell lineage, and quantified the distribution of microglial cells in five main brain regions (subventricular zone, svz; striatum, st; white matter, wm; cortex, ctx; and cerebellum, cere). The graph in Figure 3 represents the average number of cells per each area after counting 12-16 consecutive sections separated 140 μm apart. Once again, the intensity of the signal and the distribution of the microglial cells did not indicate that there is any abnormality with the development of this cell type in PCCA-/- animals.

### Neural markers are not altered in adult PCCA-/- hypomorphic mice

During the course of this study, a second mouse model for PCCA deficiency became available from the laboratory of Dr. Michael Barry. This mouse is a hypomorphic mouse developed in the PCCA-/- background. It retains about 2% enzyme activity, exhibits growth delays and some of the phenotypic traits of PA patients like elevated proprionyl-carnitine, methylcitrate, ammonia and glycine and signs of cardiomyopathy (Guenzel et al., 2013, Tamayo et al., 2020). An obvious advantage of these mice versus the Miyazaki PA mouse model that we used for the above experiments is that they reach adulthood and live much longer, being thus amenable to study brain phenotypes *in situ*. Our studies in the adult PCCA-/- mice (A138T mutation) include analysis in two different cohorts of mice with, at least 2 homozygotes and 2-3 heterozygotes and/or WT littermates. Before analysis, mice were evaluated for their serum ammonia levels as described in Guenzel et al. (2013). All mutant animals chosen for this study presented high levels of ammonia of >1000 ug/dL.

#### Neuronal development in adult PA mice

The adult PCCA-/- hypomorphic mice exhibit growth delays and a somewhat smaller size, consistent with prior reports (Guenzel et al., 2013). However, we did not observe any significant changes in overall brain development, similar to our results in the PCCA-/- neonatal mice. Both neuronal body markers (NeuN, **Figure 4**) and axonal markers (MAP2 and TUJ1, not shown) in the KO mice appear similar to the WT or the heterozygote animals. We did observe a marginal increase in the size of the cortical area with concomitant decrease in the size of the subcortical white matter of some KO mice (Figure 4C). However, this alteration was inconsistent among the animals analyzed. In addition, staining of adult brain sections to detect MBP (myelin basic protein) did not show any evidence of hypomyelination in the KO mice either.

**Figure 4.**
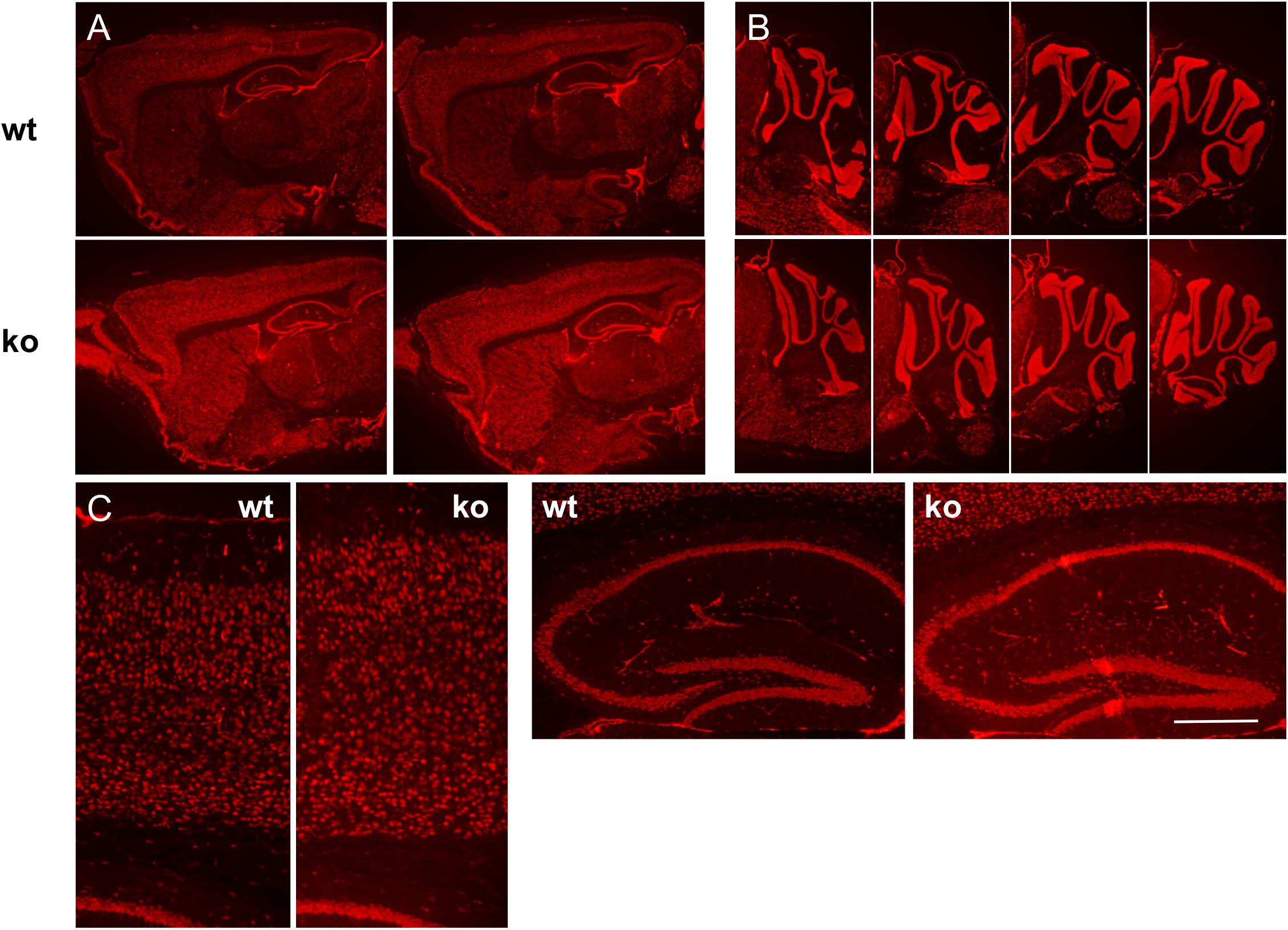
Brain development in PCCA -/- adult mice is overall normal. NeuN staining of different brain sections to analyze neuronal distribution in PCCA-/- brains of 8 week-old mice. No gross abnormalities are observed in sections obtained 140 μm apart in cortex, hippocampus, striatum (A) or cerebellum (B). Higher magnification images reveal normal hippocampus but slightly denser cortex and smaller white matter layer (C). Scale bar, 500 μm (A, B), 100 μm (C).

#### Levels of Astrocytic markers in relation to hyperammonemia

The adult PA mouse allowed us to directly examine the astrocyte population *in vivo*, where all the astrocytic cellular interactions are preserved. Surprisingly, the degree of GFAP expression (to evaluate astrogliosis) in PCCA-/- mice was also indistinguishable from what we observed in WT/heterozygous animals (**Figure 5)**. This was unexpected given that the PA mice experience hyperammonemia similar to what has been described in human patients. Similarly, the immunoreactivity levels for glutamine synthetase (GS), the main ammonia-detoxifying enzyme in brain, were also unaltered in the PCCA-/- mice. This observation is in sharp contrast to what we found in another mouse model of hyperammonemia: the ornithine transcarbamylase deficient mice (OTC-spf), where both GFAP immunoreactivity and GS levels are significantly increased.

**Figure 5.**
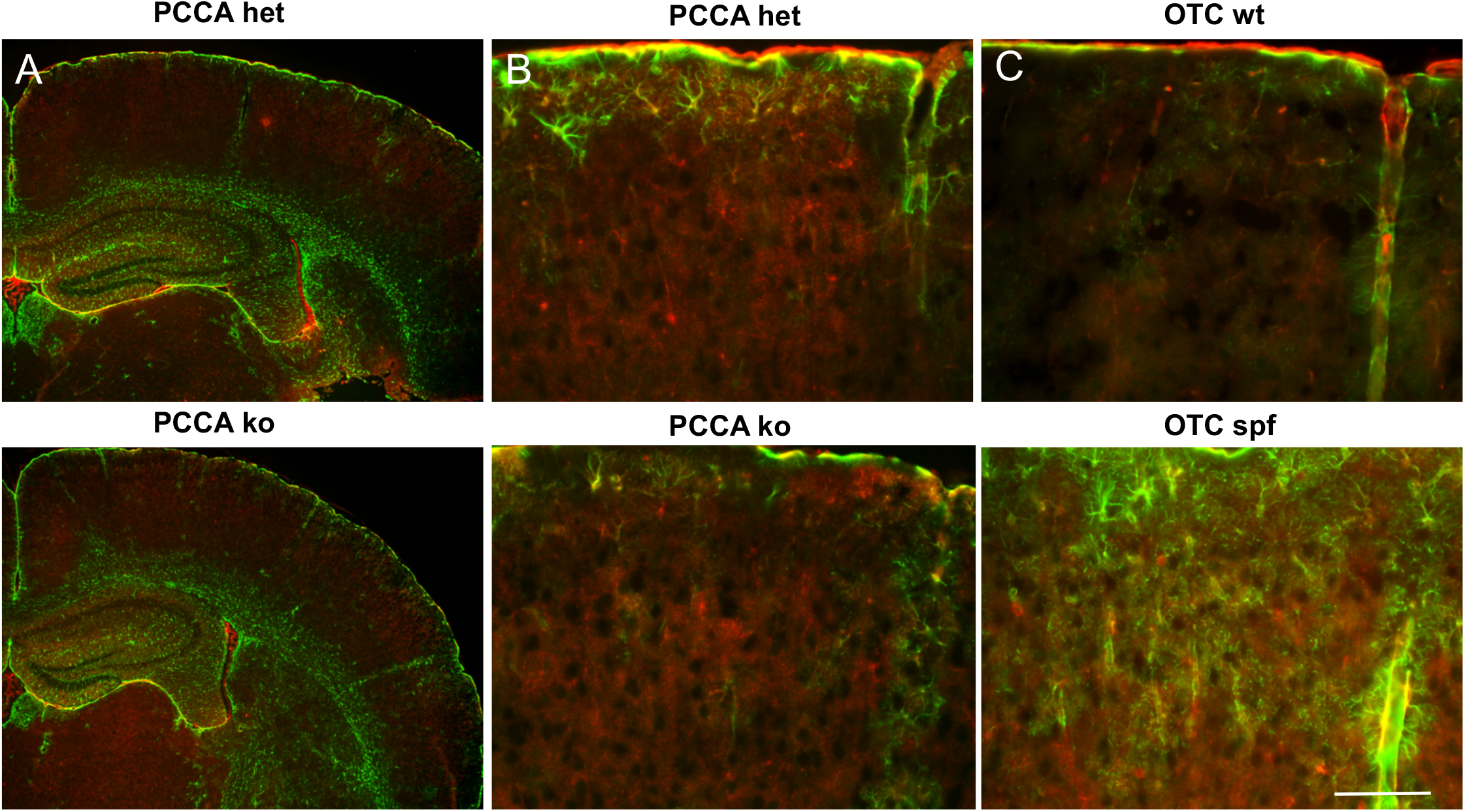
Hyperammonemia promotes reactive gliosis in OTC mice but not in PA mice. GFAP (green) and GS (red) immunoreactivity does not change in adult PCCA ko mice compared to PCCA het mice (A, B) but is greatly increased in the OTC-spf mice compared to wild-type (OTC-wt) (C). Scale bar, 250 μm (A), 50 μm (B,C).

#### Inflammation in adult PA mice

Analysis of the pattern of immunoreactivity of the marker Iba1, which stains microglia, the immune cells of the brain, shows no clear signs of reactive response in any of the areas analyzed in PCCA-/- mice, although one animal of the litter showed clear microgliosis that was most prominent in the cortex (**Figure 6**). An unremarkable presence of significant microgliosis is consistent with what we have observed in the other hyperammonemia mice in the lab, the OTC-spf mice, although a great deal of variability was again observed among different animals.

**Figure 6.**
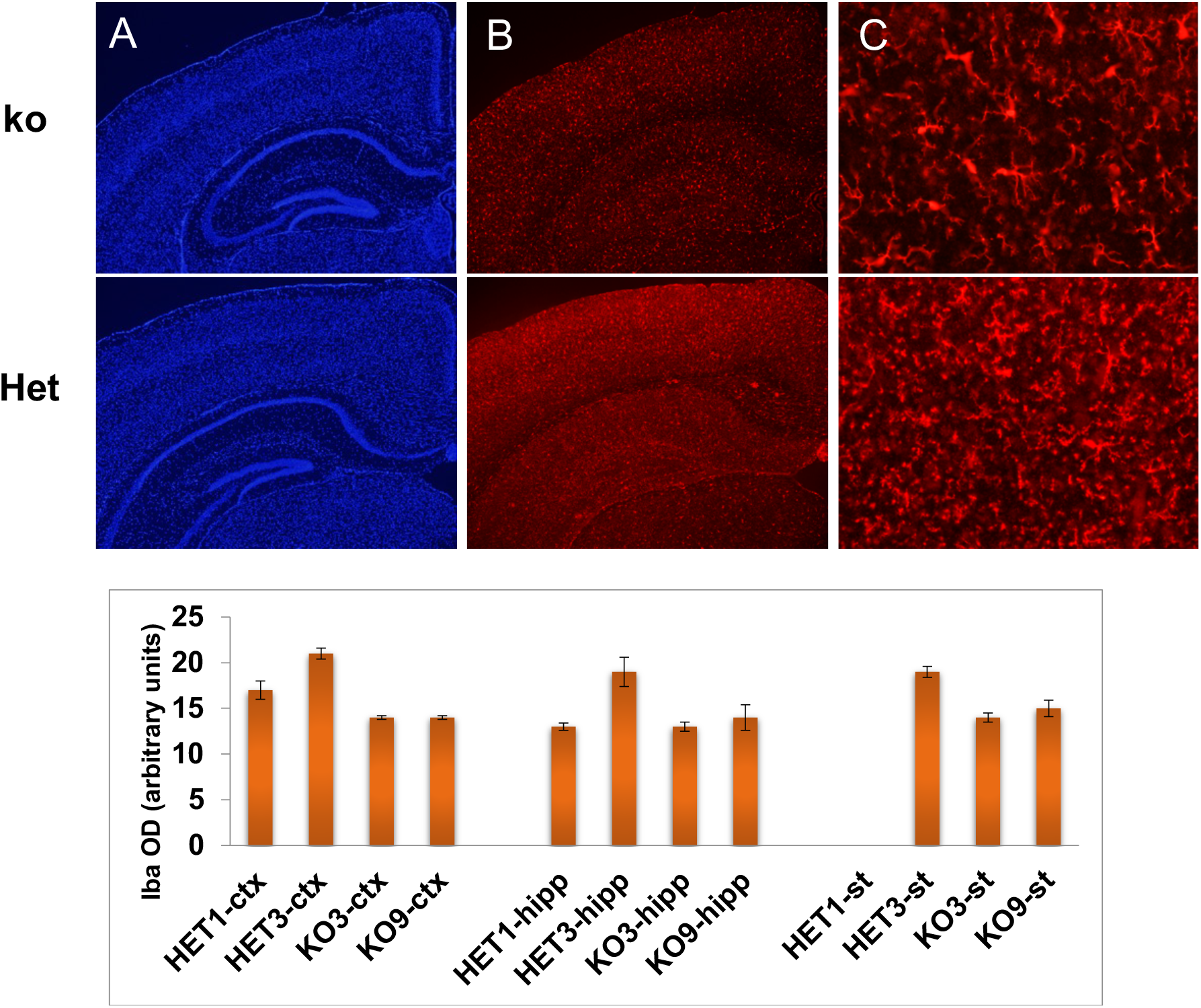
Variable signs of microgliosis in PCCA-/- mice. Iba positive cells retract their processes in the ko3 brain, an indication of microglia response to injury. However, there is high variability in the degree of microgliosis in both wt/het and ko animals (bottom graph) being the cortex the area with most microgliosis. Scale bar, Blue (DAPI), Red (Iba).

### Astrocytes respond to therapies that improve mitochondrial morphology and function

Supplementation with L-carnitine has proven effective in patients affected with PA because of its capacity to conjugate and eliminate toxic acyl-CoA derivatives (Roe et al., 1984). To indirectly study whether astrocytes could be a direct target of carnitine, we tested the effects of L-carnitine in cultured wt cortical astrocytes after exposure to hydrogen peroxide, which causes oxidative stress and cell death. Astrocytes exposed to sub-lethal doses of hydrogen peroxide survive significantly better after a 2-hour incubation with L-carnitine (**Figure 7B**). Also, mitochondria in L-carnitine-treated astrocytes are present in larger aggregates (**Figure 7A**), indicative of a more efficient respiratory chain (Mitra et al., 2009).

**Figure 7.**
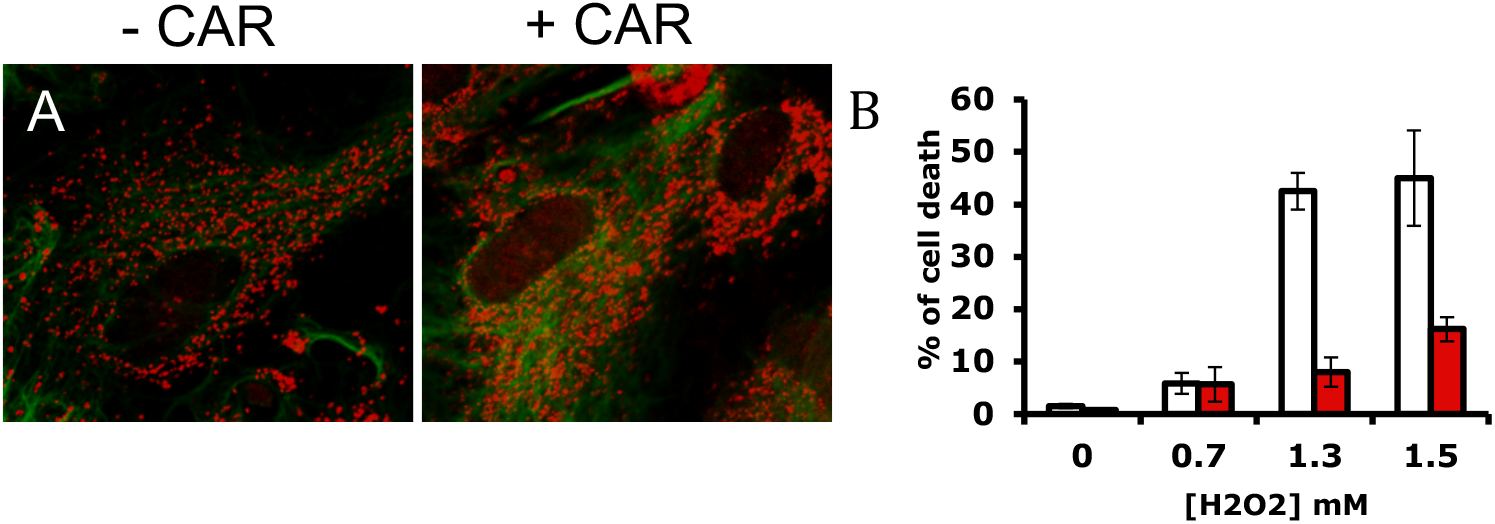
A). Mitochondria morphology in astrocytes treated with carnitine for 2 hours after 1.5 mM hydrogen peroxide (red, mitochondria; green, GFAP). B). Quantification of the effect of carnitine on astrocytic cell death after hydrogen peroxide treatment (white, control; red, carnitine-treated). CAR, carnitine.

## DISCUSSION

### Astrocytes are not a major component in the pathology of PA

The main conclusion from our study is that, surprisingly, astrocytes do not seem to be a direct component in the pathology of PA, at least under the conditions we studied.

First, in our experiments of transcriptome analysis, we found no difference between astrocytes derived from KO or from WT animals. This set of results was very surprising as we expected astrocytes to be somehow impaired by the absence of PCCA, given their wide involvement in brain metabolic support.

Second, we did not find any signs of differential levels in our study of astrocytic and neuronal markers of adult PCCA-/- brains. It was also surprising not to find signs of reactive astrogliosis, even when the adult mouse model shows signs of hyperammonemia similar to the other hyperammonemia model that we have studied in the lab, the OTC-spf mouse and where the astrocytes are highly responsive.

The fact that GS was similarly not increased in the PA mice supports clinical observations of equal or somewhat low glutamine levels in PA patients, as opposed to the urea cycle disorders, for example, that also show chronic hyperammonemia in brain (reviewed in Matsumoto et al., 2019). It is assumed that this low glutamine is due to the depletion of the glutamate-glutamine pool because of a deficient TCA cycle caused by PCCA deficiency. Another plausible explanation, however, could be that ammonia is actually detoxified by another route in the PA brain, which does not involve GS activity.

### PCCA deficiency does not cause significant changes in brain development or significant neuronal loss

From the DAPI nuclear staining of the mice brain, that can detect pyknotic (dying) nuclei, we did not observe any traces of enhanced cell death in any of our PCCA-/- animals. Consistent with this result, we did not find significant changes in the organization of the neuronal layers of the brain or cerebellum or changes in the myelination pattern. This is important because of the current clinical trend of using liver transplantation to restore normal metabolite levels in blood of PA patients. If indeed, the brain of PA patients is not affected by intrinsic toxicity of PCCA deficiency, we should not see an increased risk of metabolic stroke in patients with good metabolic control after liver transplantation. A long-term observation of liver-transplanted patients is necessary to confirm if this is the case.

It is possible that PCCA is not relevant in culture conditions where all nutrients are in excess and that, similar to what is often observed in patients affected by PA, the effects of PCCA deficiency may be only noticeable under stress conditions. Another possibility is that neurons, and not astrocytes, are the only cells affected by PCCA deficiency. This would imply that, in order to characterize the involvement of astrocytes in the pathology of PA, we would need to study them *in situ*, where all the cell interactions are maintained intact.

### Astrocytes respond to treatments that improve mitochondrial morphology

An important observation of this study was the effect of carnitine in mitochondrial functioning and survival in cultured astrocytes. Carnitine is one of the few medications proven effective in the treatment of PA, by being and adjuvant in the elimination of toxic metabolites. This could indicate a positive effect of carnitine at the level of mitochondrial machinery and improved survival in astrocytes although more studies on mitochondrial function would be needed to prove this point.

### The PA PCCA-/- mouse as model to study brain damage in PA

From the results obtained from our gene expression study *in vitro*, in the PA mice, astrocytes do not seem to be the primary cell type involved in responding to the damage caused by PCCA deficiency, even though they are the primary cell in contact with the vasculature (with high circulating ammonia levels) and are involved in metabolic stability in brain. However, changes in the vasculature and deficits in glucose utilization may play an important role in the brain damaged observed in PA patients.

At this point and given the lack of observable phenotype obtained from our studies in the PCCA-/- mice or the hypomorphic mouse, it is critical to validate the PA adult mouse model as a tool to study brain pathology in PA. Our data has proven so far inconclusive in this aspect. Studies where the brain is subject to metabolic stress may be needed to verify that similar basal ganglia damage is obtained in this mouse model. A similar issue arose in studies done in the mouse model for Glutaric Aciduria, another organic acidemia with mutations in the glutaryl-CoA dehydrogenase (GCDH) and disrupted catabolism of lysine and tryptophan. Whereas the GCDH-/- initial mouse model resulted in essentially healthy animals, exposure to a Lysine enriched diet produced a clear neuropathology of acute bilateral striatal damage similar to the one observed in young children affected by this disease (Zinnanti et al., 2006) So far, we have not observed any signs of metabolic stroke in the brains of the PA animals.

Once brain pathology similar to what is found in patients is achieved in the KO mice or rats, we will be able to test potential therapeutic compounds that improve brain health in PA.

## ACKNOWLEDGMENTS

This work has been supported by the NIH Bridges to the Baccalaureate Program (JZ). The authors would like to thank the Propionic Acidemia Foundation for its relentless mission towards finding a cure for patients with propionic acidemia.

